# The distribution of the Lansing Effect across animal species

**DOI:** 10.1101/2022.04.27.489689

**Authors:** Edward Ivimey-Cook, Sarema Shorr, Jacob Moorad

## Abstract

Maternal senescence is a reduction in individual performance associated with an increase in its mother’s age at conception. When manifested on adult lifespan, this is known as the ‘Lansing Effect’. Single-species studies report both maternal age-related increases and decreases in adult lifespan, but no comprehensive review of the literature has yet determined if the Lansing Effect is a widespread phenomenon. To address this knowledge gap, we performed a meta-analysis of maternal aging rates taken from all available published studies. We recovered 74 estimates from 20 studies representing 14 species. All studies taken together suggest a propensity for a Lansing Effect with an estimated average effect of maternal age on adult lifespan of between -22% to -17% (the lifespan response to an increase in maternal age expressed in the same time units), depending upon our specific choice of model. We failed to find a significant effect of animal class or insect order, but given the oversampling of insect species in the published literature and the paucity of vertebrate studies, we infer that only rotifers and insects demonstrate a tendency for expressing the phenomenon.

## Introduction

Senescence is the association between increased age and the deterioration of organismal function as manifested upon key components of fitness such as survival and fertility (termed actuarial and reproductive senescence, accordingly), but it can also be observed in many other traits that may affect survival and reproduction (functional senescence). Most discussions of senescence relate the ages and phenotypes of the same individuals, but recent attention has begun to focus on the social effects of aging, or how the age of one individual affects the outcomes of one or more others. Relevant social interactions can involve the ages of grandmothers (Hawkes 2003, Moorad and Walling 2017), mothers (Rogers 1993), siblings (Hamilton 1966), and even residents of the same population (Ronce and Promislow 2010), but most studies involve maternal age effects because maternal-offspring relationships are generally considered to be the most important social interactions across a great diversity of plant and animal species (Mousseau and Fox 1998). Most of these studies focus upon two effects of maternal age: pre-adult survival and lifespan.

The most closely studied of these manifestations of aging has been pre-adult survival, likely because this trait is understood to be a key component of fitness and an otherwise important trait of interest to many fields of biology. Evolutionary genetic models predict widespread maternal senescence for this trait, especially at older maternal ages (Moorad and Nussey 2016), and a recent survey of the published literature finds high prevalence of this sort of aging across all well-studied animal groups, with birds representing a notable exception (Ivimey-Cook and Moorad 2020). The second well-studied aspect of maternal aging is its manifestation upon offspring adult lifespan (which together with juvenile survival describes total lifespan). A decrease in longevity associated with an increase in maternal age at birth is known as the ‘Lansing Effect’ in recognition of Albert Lansing’s observations of the phenomenon in parthenogenic rotifers (Lansing 1947). Follow-up studies failed to replicate Lansing’s results in rotifers (Comfort 1953, King 1983), but many other studies have found mixed evidence for a Lansing effect in numerous species, including humans (Galipaud and Kokko 2020, Monaghan et al. 2020).

While awareness of the Lansing Effect appears to be high (Monaghan et al. 2020), the study of maternal age effects on offspring lifespan lags behind study of pre-adult survival in two important respects. First, evolutionary biology lacks a formal predictive genetic model of the phenomenon. However, the need for this is currently unclear owing to the second knowledge gap, which is that our current understanding of the prevalence of the Lansing Effect is only anecdotal: we lack a rigorous synthesis of the published literature that can summarize its prevalence and magnitude across studies and species. To address this concern, we have undertaken a meta-analytic review of evidence for the Lansing Effect across published studies. Our primary focus is to determine if the manifestation of a Lansing Effect is a general tendency across animal species. A secondary goal is to identify predictors for maternal age effects on lifespan, such as environment, offspring sex, and phylogeny.

## Methods

This meta-analysis followed the Preferred Reporting Items for Systematic Reviews and Meta-Analyses (PRISMA) guidelines (Figure S1). We searched for relevant published studies using the databases Web of Science (Carloni 2018) and Scopus (Baas et al. 2020) and the search terms provided in Supplemental Table S1 between February - October 2021. Studies were screened using the Rayyan web-based application (Ouzzani et al. 2016) and included in our study if they presented the following:

1. estimates derived from original data;
2. average lifespan data of individuals born to mothers of at least two different specified ages;
3. estimates of standard deviations (SDs) and/or standard errors about the means (SEs) corresponding to each maternal age; and
4. sample sizes for offspring at each maternal age class.

Data corresponding to requirements 1-4 above were extracted from the main text, supplementary information, figures, and tables of the selected studies. In cases where data were shown only in figures, the package *metaDigitise v1.01* (Pick et al. 2019) was used within *R v4.1.0* (Team 2021) to extract relevant information. Summary statistics are provided in Supplemental Table S2. From these data, we extracted or estimated the slopes of the least-squares regression of offspring lifespan on maternal age (and their standard errors) using four different methods described in descending order of preference.

1. If direct estimates of linear maternal age effects (with associated SEs) were unfettered by estimates of higher order contributions of age or another variable, the slopes were taken directly from the source paper. However, this occurred only once (Ivimey-Cook and Moorad 2020) Some studies only fit polynomial functions or other higher order interactions involving maternal ages. As we were interested only in the linear effects of age, we ignored these results and derived slopes using other methods.
2. If the raw data were provided, slopes were estimated by fitting a simple linear model that regressed offspring lifespan against maternal age class. We did this for four studies (Dowling et al. 2014, Bouwhuis et al. 2015, Lind et al. 2015, Angell et al. 2022). Where possible, and if the model converged, appropriate random effects (for instance maternal ID) were incorporated to provide a clearer estimate of maternal aging.
3. For all other cases, we assumed that observations of offspring lifespan were independent of one another and distributed normally. We applied an optimization procedure to estimate the slopes of least-squares regressions and their associated SE using maternal-age-specific means and SEs taken from the publications (see Supplement for further explanation).

Our analyses took the form of three multi-level meta-analysis models, each applied to two overlapping age ranges. Species, study, and replicate ID were fit as nested random effects in each. The first model used slope estimates to test for an overall effect size of maternal age on lifespan. The second model extended the first to evaluate and to correct for the presence and effects of publication bias. Following recommendations by Nakagawa et al (2022), publication bias was examined statistically by regressing point estimates against their respective SEs. An additional moderator of mean-centred publication year was added to the above model to test and account for time-lag bias (an association between date of publication and effect size). The intercept fit to this model is interpreted as the overall average slope that is ideally unbiased by time lag and publication bias. Noble et al (2017) suggest that the comparison between this intercept and one derived from a model that does not include these moderators can be interpreted as a test for result robustness. Monaghan et al (2020) has recently pointed out that highly concave aging trajectories may make it difficult to resolve Lansing Effects if offspring lifespan from young mothers are included in simple linear regressions. For this reason, we apply all models to data from two ranges of maternal ages. *All* contains the complete data extracted from each study. *Old* is comprised of data from maternal ages determined by one of two methods, depending upon the study and species. For the two vertebrates species, *old* ages are those that exceed the generation time *T*, or the average age of mothers at birth (estimates for *T* were taken from Felsenstein (1971) for Homo sapiens and Sæther et al. (2013) for *Sterna hirundo*). Whilst no theory yet exists to make general predictions relating to the age of onset of a Lansing Effect, evolutionary models of maternal age effects predict that *T* should predict the age of onset for maternal senescence on juvenile survival (Moorad and Nussey 2016). *Old* mothers were restricted to the last two ages for invertebrate species, as estimates of *T* were either unavailable or highly sensitive to environmental conditions, particularly temperature (Cui et al. 2018). All ages are considered *old* in studies that considered only two ages classes; we assume that the relevant published experimental designs have chosen informative ages with which to investigate the phenomenon.

The third model was used to evaluate the influence of moderator variables. These factors were:

1. The species ‘group’, defined as Order for insects and Class for non-insects;
2. Paternal-age-controlled (PAC: yes or no);
3. offspring sex (male, female, or pooled).

The effect of laboratory or natural environment was initially considered as a moderator, but this was rejected due to the low number of slopes that were able to be estimated from field studies (Table 1) and the large prevalence of certain classes only present in the laboratory (e.g., insects - see Table 1). As insect species were far better represented in the literature than non-insect species (10 vs 4 species), the former were grouped at a lower taxonomic rank to allow for the finer-scaled comparisons. This model was applied independently to both age ranges and fit with and without bias moderators.

**Table 1.** Joint distribution of major species groups and laboratory/field environments described by all extracted studies.

|  | Laboratory | Field |
| --- | --- | --- |
| **Birds** | - | $k = 2$, $n = 1$, $s = 1$ |
| **Mammals** | - | $k = 12$, $n = 3$, $s = 1$ |
| **Insects** | $k = 51$, $n = 13$, $s = 9$ | $k = 2$, $n = 1$, $s = 1$ |
| **Other Invertebrates** | $k = 7$, $n = 2$, $s = 2$ | - |

All aforementioned factors were fit into our model as fixed effects with species, study, and replicate number fit as nested random effects to account for non-independence of slopes (Nakagawa and Santos 2012; Nakagawa et al. 2021b). All subsequent analyses and visualisations were carried out in *R v4.1.0* (R Core Team 2021), using the *metafor v3.0.2* (Viechtbauer and Viechtbauer 2015), *ggplot2 v3.3.5* (Wickham 2009), *emmeans v1.7.1-1* (Lenth and Lenth 2018) *and orchaRd v0.0.0.9* (Nakagawa et al. 2021a) packages.

## Results

We extracted or estimated a total of 74 slopes (*k*) derived from 20 published studies (*n*) of 14 species (*s*). Of these, 52 estimates (70.3%) came from studies with more than two maternal age classes. All derived estimates of the slopes (and SEs) are provided in the Supplemental Section (Table S2). 16 estimates came from field studies (*n* = 5, *s* = 3), and the remaining 58 effect sizes came from laboratory studies (*n* = 15, *s* = 11) (Table 1). Whilst the dominant field organism was *Homo sapiens* (*k* = 12, *n* = 3), the most studied species in the laboratory was *Drosophila melanogaster* (*k* = 28, *n* = 4). As expected, insect species dominated the laboratory environment (*k* = 51/58), and a study of the antler fly (*Protopiophila litigata*, Angell et al. 2022) provided the sole estimate derived from a natural insect system (Table 1). Strong associations between taxonomy and environments, namely the observation that insects are rarely studied in the field and mammals and birds rarely studied in the laboratory, resemble those reported in a recent review of the relationship between maternal age and early offspring survival (Ivimey-Cook & Moorad, 2020).

We fit all extracted slopes to a progression of three models (Models 1 – 3) using all available maternal ages (Table 2 – All). Model 1 is the intercept-only model; this indicates a clearly negative value for the estimates (−0.22) with 95% confidence intervals that do not overlap zero (95%CI = -0.33, -0.12; *p* = 0.00005). Differently put, offspring lifespan is observed to decline by 22% of the increase in maternal age. Model 2 adds estimates for the effects of publication bias and time lag. Neither of these effects have a statistically significant effect on the slopes (*p* = 0.326, 0.766, accordingly), but including these moderators reduces the intercept from -0.22 to -0.17, and the confidence intervals associated with the intercept widen but still do not overlap zero (95%CI = -0.32, -0.007). However, these estimates may be conservative: Stanley and Doucouliagos (2014) report that the estimate of the intercept will be a downwardly biased indicator of the magnitude of the true mean, which, in this case, indicates that the true mean effect of maternal age (once publication and time lag biases are accounted for) is less than -0.17.

**Table 2.**
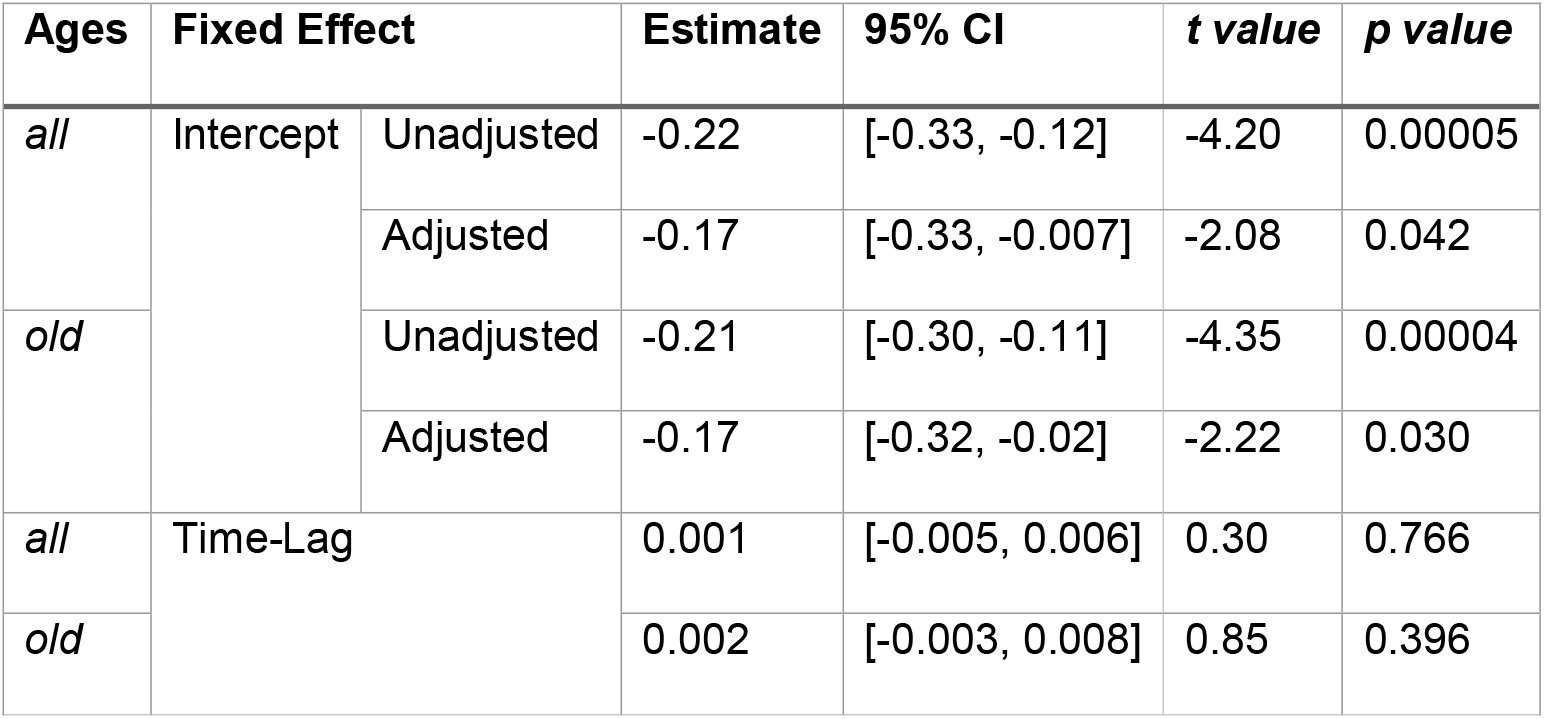
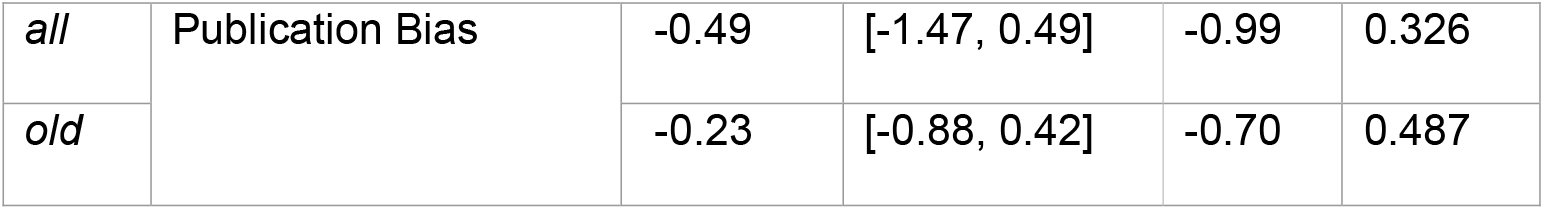
Meta-analytic mean slope unadjusted (Model 1) and adjusted (Model 2) for publication bias (standard error) and time-lag bias (mean-centred year) with corresponding 95% confidence intervals, *t*, and *p* values. Estimates for the effects of bias derived from Model 2.

Focusing our analysis only on the last two maternal age classes (Table 2 – *old*) makes very little difference in the estimates other than to reduce the unadjusted age-related decline in offspring lifespan from 21% to 17% and to narrow the confidence interval associated with the adjusted estimate enough for the point estimate to be significant at the α = 0.05 level (−0.32, -0.02).

Next, we considered the effects of species group, PAC, and offspring sex (Model 3). We fit this model with and without time lag and publication bias moderators and for all ages and the terminal interval (the model was fit four different ways). As before, we find no significant effect of the bias moderators, and the choice of which ages to include has little effect on the results. Table 3 summarizes results from the version of the model without bias moderators and with *All* age classes; other results are provided in the Supplement (Table S3). A comparison of marginal means for all groups (Fig. 1) finds the strongest evidence for a Lansing Effect in Orthopterans, Dipterans, Coleopterans, and rotifers. Hemipterans show a similar pattern when all ages are considered, but the Lansing Effect disappears when these are analysed over only the terminal ages. Neither nematodes, mammals, nor birds appear to present tendencies for Lansing Effects. However, it should be emphasized that no differences among these groups are statistically significant in any version of the model. Considering both the estimates and the sample sizes of slopes taken from the various animal groups, it appears that the overall support for a general Lansing Effect may be driven by an oversampling of insects, where there appears also to be the greatest prevalence of negative slopes. A visual comparison of these marginal means found no evidence to suggest that focusing only upon *old* ages would increase the statistical evidence for a tendency towards a Lansing Effect.

**Table 3.**
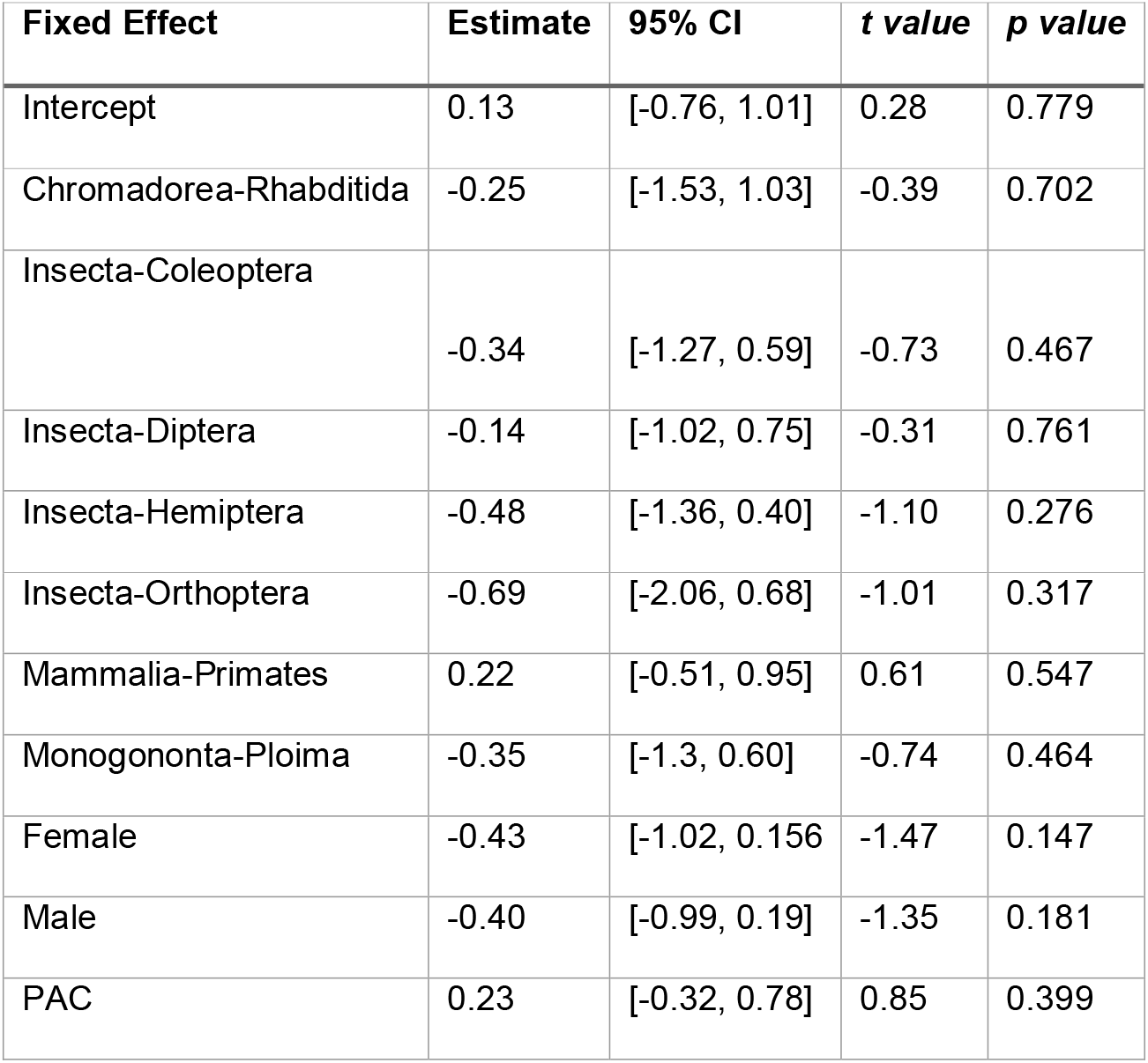
Full model output from a multi-level mixed effect model across all species with moderator variables acting on slopes. Note that the reference categories are Aves, mixed offspring sex, and uncontrolled mate age; the Intercept is the estimated mean for their combination. Effects are shown unadjusted and adjusted for publication and time lag bias.

**Figure 1.**
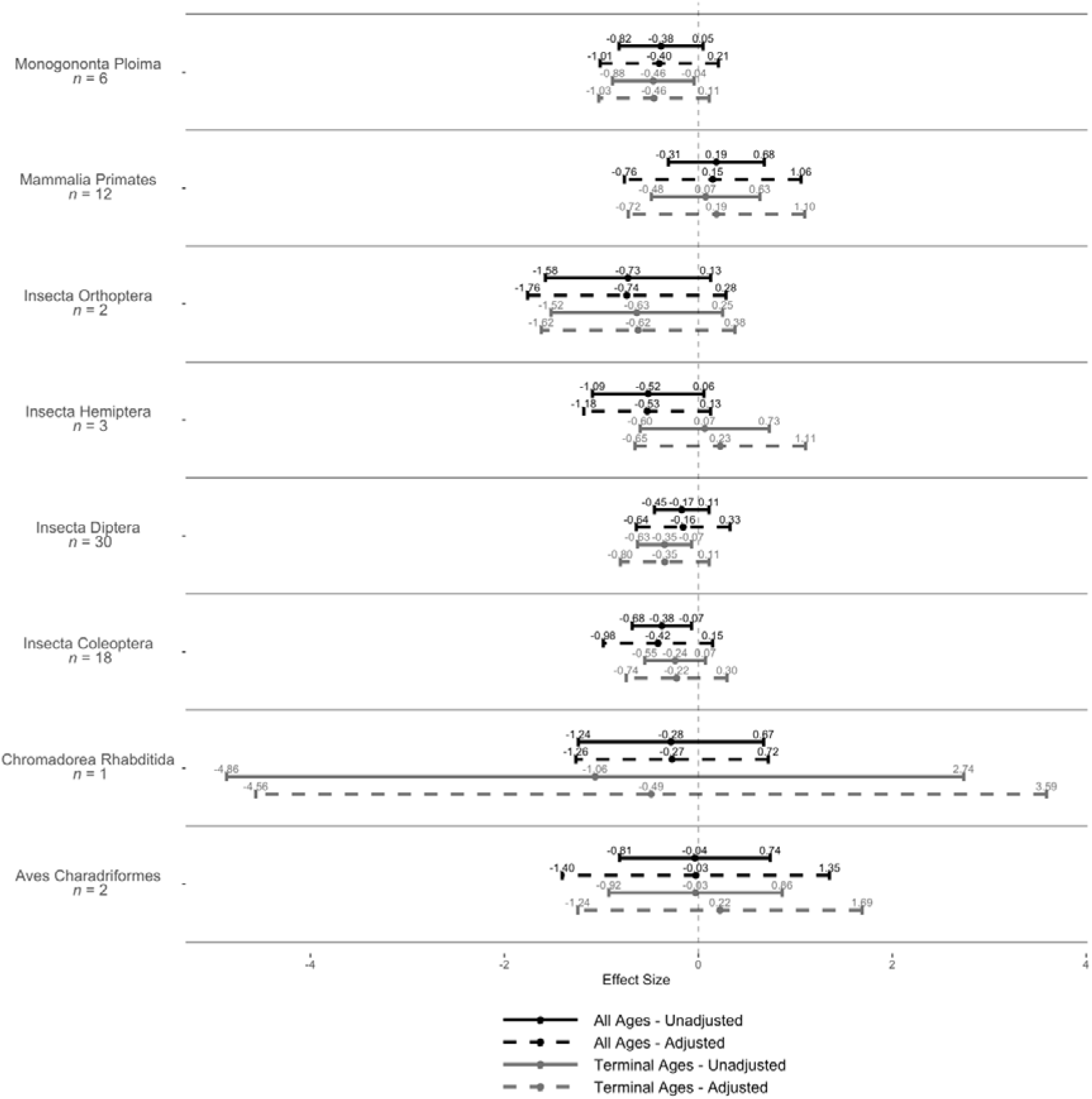
The marginal means of slope estimates for species groups (*n* = species) with corresponding 95% confidence intervals.

Controlling mate age (PAC) appears to diminish the strength of the Lansing Effect, but that effect is not statistically significant (*p* = 0.399). We might expect this direction of effect if parental ages are positively assorted *and* if a Lansing-like Effect acts through paternal age, but we lack the means to test this suggestion here. Finally, we note that the Lansing Effect appears to manifest nearly identically upon females and males and more strongly on specific sexes than on mixed-sex offspring. We have no explanation for this pattern except to suggest, with no evidence, that experimental designs that pool sexes when reporting lifespan differ in other respects from those that segregate by sex.

## Discussion

A Lansing Effect could be said to exist in any single study that finds a negative association between maternal age and offspring longevity, but a multi-study view can provide new evolutionary or ecological insights, such as a characterization of the central tendency of the phenomenon or the identification of modifiers that may provide new understanding of the causes of the Lansing Effect. The primary goal of this study was to determine if the sum of published data was sufficient to detect a tendency across animal populations to exhibit Lansing Effects. Here we were successful: we found clear evidence that such a tendency exists: a unit increase in maternal age translates to a decrease in offspring lifespan of 16-22% of that unit. However, that negative tendency appears to be driven by observations of insects and, to a lesser degree, rotifers, which are oversampled compared to other animal groups that offer little to no tendencies themselves.

Neither bird, mammal, nor nematode species demonstrated the statistically significant deleterious maternal age effects on offspring lifespan that are consistent with a general Lansing Effect. In fact, birds and humans exhibited near-zero effect sizes. This is surprising given that the conspicuous maternal care provided in these species should present more opportunities for deleterious effects of increased age to manifest. However, it could be that post-natal mechanisms of maternal care are more amenable to improvement with age owing to the accumulation of experience, and this effect mitigates or even overwhelms senescence for pre-natal maternal inputs into offspring survival. This sort of conflict between aging sources has been suggested as important for understanding juvenile mortality in seabirds (Aubry et al. 2009, Aubry et al. 2011, Froy et al. 2013). Finally, we note that post-natal parental care has been suggested to buffer against the deleterious effects of the environment (Schroeder et al. 2012, Pilakouta and Smiseth 2016, Grew et al. 2019). If true, then it may then seem logical to expect that animals such as birds and mammals should have lower rates of maternal senescence if one were to consider old maternal age as a poor environment. However, this argument neglects that possibility that buffering ability might also senesce, and this would serve to reinforce the deleterious effects of increased maternal age in systems with maternal care (see Moorad and Ravindran (2022) for a fuller discussion of buffering in the context of the evolution of aging).

Clearly, we need far more relevant studies of vertebrate species featuring varying degrees of post-natal care if we wish to understand the prevalence (let alone the causes) of the maternal age effects on lifespan in birds and mammals. Reproductive experience (or parity) should be controlled for experimentally or statistically, and further attention should also focus on species and systems that can allow us to disentangle taxonomy and environment, for example by studying vertebrates in the laboratory or insects (or other invertebrates) in the wild.

Our results correspond very roughly to those of a recent review of maternal age effects on juvenile survival (Ivimey Cook and Moorad 2020) in the sense that the most negative effects of age appear to manifest in invertebrates and the most positive effects in birds. Mammals are intermediate in both meta-analyses, but whereas the previous study had sufficient statistical power to bound grouped effect sizes away from each other (e.g., invertebrates and mammals had significantly negative effects, and birds had significantly positive effects). We note that the current study found considerably less relevant metadata than the juvenile survival study; it is much more difficult to measure lifespan than juvenile survival in most laboratory and wild animal populations, and this difference is likely reflected in the quantity and precision of estimates that are available from the published literature. The current study extracted 74 estimates from 14 species, with a median number of offspring for each estimate of 96. This compares poorly with the Ivimey-Cook and Moorad (2020) study, with 273 estimates from 97 animal species and a median number of offspring per estimate of 234. This discrepancy underscores the need for more studies of the Lansing Effect to be done on more species.

Although the current study failed to detect statistically significant differences among species groups, our observations of a general tendency towards a Lansing Effect and similar rankings of species groups when we consider the strength of the two different manifestations of maternal senescence merit attention from evolutionary theory. Does the theory predict that age-specific selection that acts on maternal effect genes for adult lifespan invariably weaken with age similar to how we expect age-specific selection for direct effect genes to relax (Hamilton 1966, Charlesworth 1994, Moorad and Ravindran 2022)? Alternatively, does this pattern of selection have a maximum at some later maternal age, as we expect can happen for maternally-expressed genes that affect juvenile survival (Moorad and Nussey 2016)? It seems intuitive to expect that selection for a Lansing Effect should have dynamics that are more similar to the latter, and this would help explain the rough congruence between the rankings of animal group aging rates, but a proper answer to this question awaits development of the relevant evolutionary genetic theory.

Finally, we offer some recommendations for future studies on maternal senescence. First, maternal ages that clearly represent old individuals should be included in observations, and justifications for why these ages are considered as such should be provided. We advocate the use of mean generation time *T* as a useful yardstick in this effort because it defines the mean age of parents in the population, and this determines exactly what it means for a parent to be older than average. Second, experimenters should be aware that cohorts of same-age individuals change over time for reasons owing to aging (within-individual changes) and selective disappearance (among-individual change). As we are most often interested only in the former, care should be taken to decouple these two components of change to avoid biasing our interpretations of the true aging rate. This is accomplished by fitting appropriate statistical models that include some aspect of age-at-death as a modifier (van de Pol and Verhulst 2006, Nussey et al. 2011, Ivimey-Cook and Moorad 2018), and this has become common practice in studies of conventional perspectives of aging that seek to understand the association between trait values and ages of the same individuals (Nussey et al. 2011, Hayward et al. 2013, Hamalainen et al. 2014). Applied to maternal senescence, the offspring trait should be evaluated against both the mother’s age at birth and the age of her death (e.g., Nussey et al. 2006, van de Pol and Verhulst 2006, Bouwhuis et al. 2010, Schroeder et al. 2012, Ivimey-Cook and Moorad 2018, Lord et al. 2021), but this is seldom done in practice. It should be noted that only two of the estimates used in this study derive from analyses that adequately correct for selective disappearance (Ivimey-Cook and Moorad (2018) and Bouwhuis et al (2015) either provide relevant estimates in the papers or sufficient data that allowed us to derive these). Unfortunately, no other study provided the necessary information to statistically account for these effects, and there is a risk that our results for other studies are biased. The direction of bias will depend upon the nature of the relationship between age-specific maternal survival and offspring lifespan, but if we assume that mothers vary in overall quality (i.e., longer surviving mothers are also better mothers, and this is reflected in longer offspring lifespan), then we might expect that biases work to reduce the severity of the Lansing Effect. If sufficiently strong, selective disappearance may even cause the direction of aging to be reversed. In light of this, we should consider our findings to be conservative with respect to the general tendency of animal species to exhibit a Lansing Effect. It may also be that among-group differences in selective disappearance have contributed to apparent (albeit statistically non-significant) among-group differences in the presence of a Lansing Effect.

This is the first comprehensive review of maternal senescence manifested on adult lifespan. We found a general tendency for insect and rotifer species to exhibit a Lansing Effect. This supports the notion that there are transgenerational mechanisms for the inheritance of aging; this is important to our understanding of the evolution of life histories, and it may have important implications to conservation. However, it may be premature at this point to assume that a tendency towards a Lansing Effect exists in birds and mammals. This may change as more observational and experimental aging studies are performed and evolutionary theory is developed sufficiently to know whether natural selection tends to favour the evolution of a Lansing Effect in the general case.

