## Supplementary Material for "The distribution of the Lansing Effect across animal species"

**Figure S1.** PRISMA (Preferred Reporting Items for Systematic Reviews and Meta-Analyses) flow diagram showing the various steps of obtaining studies and replicates for analysis.

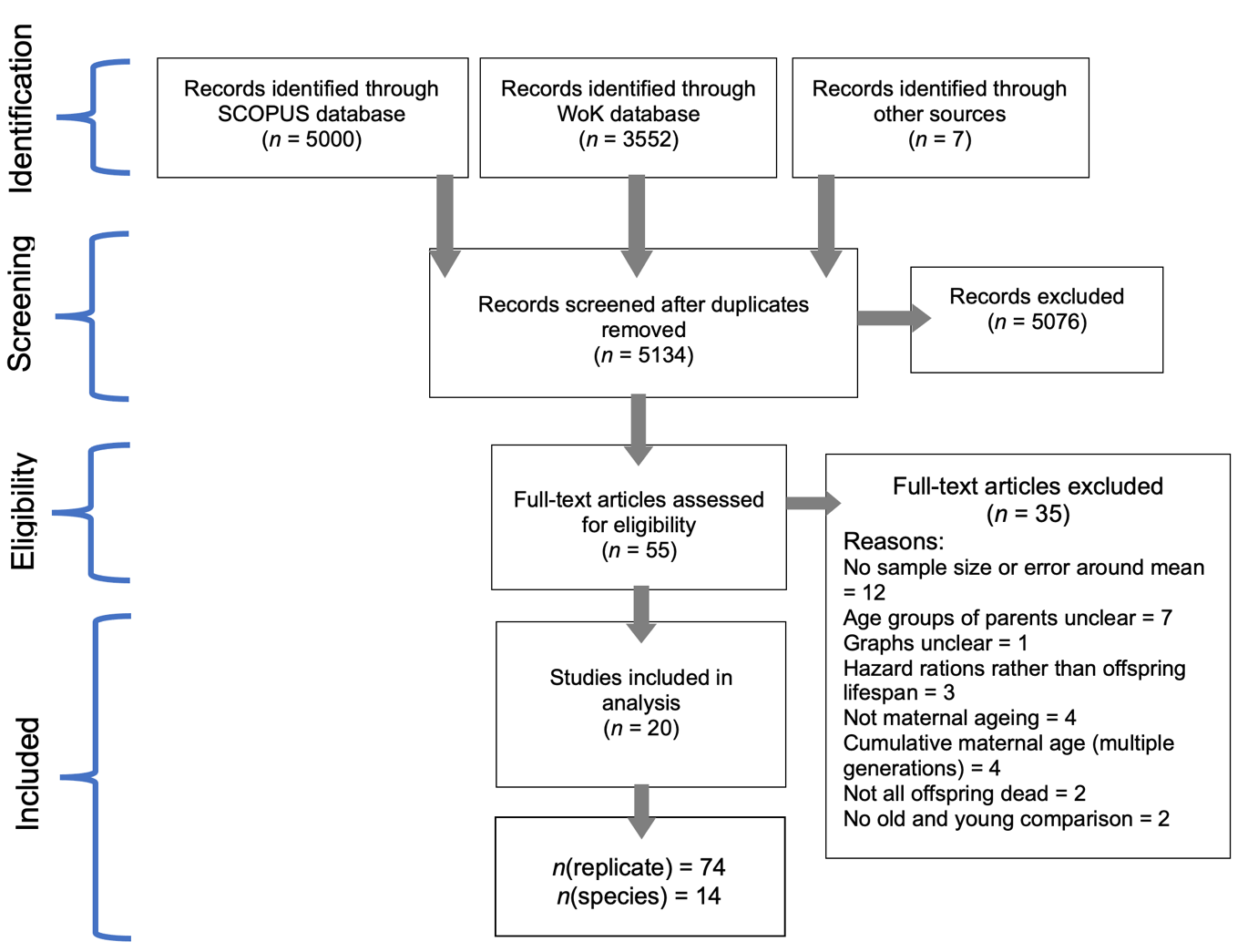

**Table S1.** Search terms used in the initial database search for both Scopus and Web of Knowledge. Asterisks (*) denote wild card terms where all permutations after the first part of the word are included.

| *Terms related to mothers* | *Terms related to age* | *Terms related to offspring* | *Terms related to longevity* | *Additional term* |
| --- | --- | --- | --- | --- |
| Mother | Age* | Offspring | Longevit* | Lansing effect |
| Maternal | Aging | Progeny | Lifespan | - |
| Female* | Senescen* | Juvenile | Life expectancy | - |
| Wom* | Old | Infant | - | - |
| Dam | - | Child* | - | - |
| - | - | Young* | - | - |

**Table S2.** Effect sizes and standard errors for all ages (All/AllSE) and terminal age classes (Terminal/TerminalSE).

| Study | Replicate | Title | Year | Class | Order | Species | Lab / Field | PAC | Offspring Sex | All | AllSE | Terminal | TerminalSE |
| --- | --- | --- | --- | --- | --- | --- | --- | --- | --- | --- | --- | --- | --- |
| 1 | 1 | Maternal age and lifespan of offspring | 1997 | Mammalia | Primates | *Homo sapiens* | Field | No | Male | 0.00017 | 0.03574 | -0.06451 | 0.05883 |
|  | 2 |  |  |  |  |  |  |  | Female | -0.07690 | 0.05240 | -0.26496 | 0.08699 |
|  | 3 |  |  |  |  |  |  |  | Male | -0.03635 | 0.05488 | -0.01588 | 0.08855 |
|  | 4 |  |  |  |  |  |  |  | Female | -0.10095 | 0.07504 | -0.32178 | 0.11953 |
|  | 5 |  |  |  |  |  |  |  | Male | 0.04924 | 0.02857 | 0.05783 | 0.04678 |
|  | 6 |  |  |  |  |  |  |  | Female | 0.02208 | 0.03382 | -0.08326 | 0.05531 |
| 2 | 1 | Parental age at conception and offspring longevity | 1997 | Mammalia | Primates | *Homo sapiens* | Field | No | Female | -0.02683 | 0.08921 | -0.17741 | 0.14408 |
|  | 2 |  |  |  |  |  |  |  | Male | -0.02082 | 0.06746 | -0.07883 | 0.10848 |
| 3 | 1 | Parental effects on offspring longevity - evidence from 17th to 19th century reproductive histories | 2004 | Mammalia | Primates | *Homo sapiens* | Field | No | Male | 0.01299 | 1.03915 | 0.17504 | 2.54325 |
|  | 2 |  |  |  |  |  |  |  | Female | -0.01571 | 1.03394 | -0.06173 | 2.45435 |
|  | 3 |  |  |  |  |  |  |  | Male | -0.15325 | 0.07522 | -0.38021 | 0.19677 |
|  | 4 |  |  |  |  |  |  |  | Female | -0.36241 | 0.08159 | -0.61078 | 0.20221 |
| 4 | 1 | Maternal caloric restriction partially rescues the deleterious effects of advanced maternal age on offspring | 2014 | Monogononta | Ploima | *Brachionus manjavacas* | Lab | Yes | Female | -0.95014 | 0.13441 | -0.40032 | 0.22280 |
|  | 2 |  |  |  |  |  |  |  | Male | -0.08374 | 0.06753 | 0.00041 | 0.15042 |
|  | 3 |  |  |  |  |  |  |  | Female | -0.49996 | 0.14411 | -0.44953 | 0.23584 |
|  | 4 |  |  |  |  |  |  |  | Male | -0.14814 | 0.06329 | -0.20018 | 0.13530 |
|  | 5 |  |  |  |  |  |  |  | Female | -0.71142 | 0.16630 | -0.55000 | 0.27914 |
|  | 6 |  |  |  |  |  |  |  | Male | -0.12473 | 0.06450 | -0.20000 | 0.13881 |
| 5 | 1 | Effects of parental age on the life cycle of the mealworm, *Tenebrio molitor* linnaeus | 1958 | Insecta | Coleoptera | *Tenebrio molitor* | Lab | No | Combined | -1.09771 | 0.33958 | -1.14954 | 0.68865 |
|  | 2 |  |  |  |  |  |  |  | Combined | -1.40918 | 0.24338 | -0.70045 | 0.50025 |
| 6 | 1 | Further studies on the relationship between parental age and the life cycle of the mealworm, *Tenebrio molitor* | 1960 | Insecta | Coleoptera | *Tenebrio molitor* | Lab | No | Combined | -0.08933 | 0.06717 | -0.15355 | 0.32486 |
|  | 2 |  |  |  |  |  |  |  | Combined | -0.07263 | 0.14462 | 0.13624 | 0.31436 |
|  | 3 |  |  |  |  |  |  |  | Combined | 1.19493 | 0.13354 | -1.54654 | 0.34037 |
|  | 4 |  |  |  |  |  |  |  | Combined | -0.43426 | 0.03721 | 0.15000 | 0.18014 |
|  | 5 |  |  |  |  |  |  |  | Combined | -0.22418 | 0.09826 | -1.17386 | 0.38162 |
|  | 6 |  |  |  |  |  |  |  | Combined | -0.35545 | 0.05614 | -0.69350 | 0.27961 |
| 7 | 1 | Evolution of differential maternal age effects on combined offspring development and longevity | 2015 | Insecta | Coleoptera | *Callosobruchus maculatus* | Lab | Yes | Female | -0.51804 | 0.06414 | -0.01474 | 0.24954 |
|  | 2 |  |  |  |  |  |  |  | Male | -0.26166 | 0.04429 | -0.25631 | 0.17436 |
|  | 3 |  |  |  |  |  |  |  | Female | -0.33233 | 0.06123 | -0.40807 | 0.24276 |
|  | 4 |  |  |  |  |  |  |  | Male | -0.31848 | 0.04328 | 0.49384 | 0.16276 |
| 8 | 1 | Strain‐specific effects of parental age on offspring in *Drosophila melanogaster* | 2019 | Insecta | Diptera | *Drosophila melanogaster* | Lab | Yes | Male | 0.21540 | 0.04906 | 0.21540 | 0.04906 |
|  | 2 |  |  |  |  |  |  |  | Female | -0.03095 | 0.02725 | -0.03095 | 0.02725 |
|  | 3 |  |  |  |  |  |  |  | Male | -0.03464 | 0.05175 | -0.03464 | 0.05175 |
|  | 4 |  |  |  |  |  |  |  | Female | 0.04986 | 0.04407 | 0.04986 | 0.04407 |
|  | 5 |  |  |  |  |  |  |  | Male | -0.03085 | 0.05815 | -0.03085 | 0.05815 |
|  | 6 |  |  |  |  |  |  |  | Female | -0.13496 | 0.04329 | -0.13496 | 0.04329 |
|  | 7 |  |  |  |  |  |  |  | Male | -0.16147 | 0.05999 | -0.16147 | 0.05999 |
|  | 8 |  |  |  |  |  |  |  | Female | -0.17694 | 0.03816 | -0.17694 | 0.03816 |
|  | 9 |  |  |  |  |  |  |  | Male | -1.36162 | 0.03668 | -1.36162 | 0.03668 |
|  | 10 |  |  |  |  |  |  |  | Female | -1.54239 | 0.03604 | -1.54239 | 0.03604 |
|  | 11 |  |  |  |  |  |  |  | Male | -0.17336 | 0.03269 | -0.17336 | 0.03269 |
|  | 12 |  |  |  |  |  |  |  | Female | -0.06241 | 0.01954 | -0.06241 | 0.01954 |
| 9 | 1 | Maternal age effects on longevity in *Drosophila melanogaster* populations of different origin | 2008 | Insecta | Diptera | *Drosophila* | Lab | Yes | Female | -1.66621 | 0.31615 | -0.13902 | 0.24840 |
|  | 2 |  |  |  |  |  |  |  | Male | -0.14879 | 0.32181 | 0.36068 | 0.38631 |
|  | 3 |  |  |  |  |  |  |  | Female | 0.53318 | 0.23920 | -0.12541 | 0.10293 |
|  | 4 |  |  |  |  |  |  |  | Male | -1.49207 | 0.29068 | 0.12019 | 0.25926 |
|  | 5 |  |  |  |  |  |  |  | Female | 0.75588 | 0.30965 | 0.06818 | 0.09799 |
|  | 6 |  |  |  |  |  |  |  | Male | -0.23109 | 0.34398 | 0.19246 | 0.25460 |
| 10 | 1 | The role of parental age effects on the evolution of aging | 2002 | Insecta | Diptera | *Drosophila melanogaster* | Lab | Yes | Female | -0.15657 | 0.04909 | -0.32340 | 0.10360 |
|  | 2 |  |  |  |  |  |  |  | Female | 0.07586 | 0.02759 | -0.54398 | 0.33252 |
|  | 3 |  |  |  |  |  |  |  | Male | -0.06553 | 0.03948 | -0.34026 | 0.13166 |
|  | 4 |  |  |  |  |  |  |  | Male | 0.09490 | 0.03232 | -0.32825 | 0.52231 |
| 11 | 1 | Maternal and paternal age effects on male antler flies: a field experiment | 2022 | Insecta | Diptera | *Protopiophila litigata* | Field | Yes | Male | 0.06818 | 0.09799 | 0.06818 | 0.09799 |
|  | 2 |  |  |  |  |  |  |  | Male | -0.12541 | 0.10293 | -0.12541 | 0.10293 |
| 12 | 1 | Maternal sexual interactions affect offspring survival and ageing | 2014 | Insecta | Diptera | *Drosophila melanogaster* | Lab | Yes | Male | -0.32825 | 0.52231 | -0.32825 | 0.52231 |
|  | 2 |  |  |  |  |  |  |  | Male | -0.13902 | 0.24840 | -0.13902 | 0.24840 |
|  | 3 |  |  |  |  |  |  |  | Male | -0.54398 | 0.33252 | -0.54398 | 0.33252 |
|  | 4 |  |  |  |  |  |  |  | Female | 0.36068 | 0.38631 | 0.36068 | 0.38631 |
|  | 5 |  |  |  |  |  |  |  | Female | 0.19246 | 0.25460 | 0.19246 | 0.25460 |
|  | 6 |  |  |  |  |  |  |  | Female | 0.12019 | 0.25926 | 0.12019 | 0.25926 |
| 13 | 1 | Disentangling pre- and postnatal maternal age effects on offspring performance in an insect with elaborate maternal care | 2018 | Insecta | Coleoptera | *Nicrophorus vespilloides* | Lab | Yes | Combined | -0.07500 | 0.11200 | 0.84830 | 1.10590 |
|  | 2 |  |  |  |  |  |  |  | Combined | 0.00100 | 0.12400 | -0.16090 | 0.77660 |
| 14 | 1 | Aging in the nematode *Caenorhabditis elegans*: Major biological and environmental factors influencing life span | 1977 | Chromadorea | Rhabditida | *C. elegans* | Lab | No | Combined | -0.12137 | 0.16767 | -1.20047 | 1.89865 |
| 15 | 1 | Heterogenous effects of father and mother age on offspring development | 2021 | Insecta | Orthoptera | *Gryllus bimaculatus* | Lab | Yes | Combined | -0.34026 | 0.13166 | -0.34026 | 0.13166 |
|  | 2 |  |  |  |  |  |  |  | Combined | -0.32340 | 0.10360 | -0.32340 | 0.10360 |
| 16 | 1 | Influence of maternal age on the fitness of progeny in the rice weevil, *Sitophilus oryzae* (Coleoptera: Curculionidae) | 2014 | Insecta | Coleoptera | *Sitophilus oryzae* | Lab | Yes | Combined | -0.03437 | 0.00935 | -0.04573 | 0.01496 |
| 17 | 1 | Effects of maternal age and environment on offspring vital rates in the oleander aphid (Hemiptera: Aphididae) | 2007 | Insecta | Hemiptera | *Aphis nerii* | Lab | No | Combined | -0.20101 | 0.06741 | 0.01310 | 0.13492 |
| 18 | 1 | Sex-specific pathways of parental age effects on offspring lifetime reproductive success in a long-lived seabird | 2015 | Aves | Charadriformes | *Sterna hirundo* | Field | No | Male | -0.32753 | 0.15237 | -0.09961 | 0.27412 |
|  | 2 |  |  |  |  |  |  |  | Female | -0.24619 | 0.22958 | -0.63018 | 0.55829 |
| 19 | 1 | Effects of parental age on the life cycle of the Southern Green Stink Bug, *Nezara viridula* L. (Heteroptera : Pentatomidae)  : Pentatomidae) | 1967 | Insecta | Hemiptera | *Nezara viridula* | Lab | No | Male | -0.87435 | 0.14945 | -0.03017 | 0.58618 |
|  | 2 |  |  |  |  |  |  | No | Female | -0.84427 | 0.15912 | -1.06079 | 0.94453 |
| 20 | 1 | Effects of temperature and parental age on the life cycle of the Dark Mealworm, *Tenebrio obscurus* Fabricius | 1960 | Insecta | Coleoptera | *Tenebrio obscurus* | Lab | No | Combined | 0.06219 | 0.07796 | 0.08447 | 0.18867 |
|  | 2 |  |  |  |  |  |  |  | Combined | -0.31940 | 0.13009 | -0.42168 | 0.20405 |
|  | 3 |  |  |  |  |  |  |  | Combined | -0.00096 | 0.03097 | -0.07066 | 0.05292 |

**Table S3.** Full model output for all moderators across all ages (Full) and terminal ages (Terminal) showing estimates, 95%CI, *t* and *p* values unadjusted and adjusted for time lag and publication bias.

| Ages | Fixed Effect | | Estimate | 95% CI | *t value* | *p value* |
| --- | --- | --- | --- | --- | --- | --- |
| Full | Intercept | Unadjusted | 0.13 | [-0.76, 1.01] | 0.28 | 0.779 |
|  |  | Adjusted | 0.21 | [-0.93, 1.36] | 0.37 | 0.714 |
| Terminal |  | Unadjusted | -0.17 | [-1.18, 0.85] | -0.33 | 0.744 |
|  |  | Adjusted | -0.01 | [-1.21, 1.19] | -0.01 | 0.989 |
| Full | Chromadorea-Rhabditida | Unadjusted | -0.48 | [-1.57, 0.6] | -0.89 | 0.379 |
|  |  | Adjusted | -0.25 | [-1.53, 1.03] | -0.39 | 0.702 |
| Terminal |  | Unadjusted | -0.25 | [-2.03, 1.54] | -0.28 | 0.784 |
|  |  | Adjusted | -1.03 | [-5.03, 2.96] | -0.52 | 0.606 |
| Full | Insecta-Coleoptera | Unadjusted | -0.71 | [-4.97, 3.55] | -0.33 | 0.740 |
|  |  | Adjusted | -0.34 | [-1.27, 0.59] | -0.73 | 0.467 |
| Terminal |  | Unadjusted | -0.39 | [-2.26, 1.48] | -0.42 | 0.678 |
|  |  | Adjusted | -0.21 | [-1.26, 0.84] | -0.40 | 0.691 |
| Full | Insecta-Diptera | Unadjusted | -0.45 | [-2.3, 1.41] | -0.48 | 0.632 |
|  |  | Adjusted | -0.14 | [-1.02, 0.75] | -0.31 | 0.761 |
| Terminal |  | Unadjusted | -0.13 | [-1.9, 1.64] | -0.15 | 0.883 |
|  |  | Adjusted | -0.32 | [-1.33, 0.69] | -0.63 | 0.531 |
| Full | Insecta-Hemiptera | Unadjusted | -0.57 | [-2.38, 1.24] | -0.63 | 0.532 |
|  |  | Adjusted | -0.48 | [-1.36, 0.39] | -1.10 | 0.276 |
| Terminal |  | Unadjusted | -0.50 | [-1.78, 0.78] | -0.78 | 0.437 |
|  |  | Adjusted | 0.10 | [-1, 1.19] | 0.17 | 0.862 |
| Full | Insecta-Orthoptera | Unadjusted | 0.00 | [-1.27, 1.28] | 0.01 | 0.996 |
|  |  | Adjusted | -0.69 | [-2.06, 0.68] | -1.01 | 0.317 |
| Terminal |  | Unadjusted | -0.71 | [-2.88, 1.46] | -0.66 | 0.514 |
|  |  | Adjusted | -0.61 | [-2.14, 0.93] | -0.79 | 0.435 |
| Full | Mammalia-Primates | Unadjusted | -0.84 | [-3.03, 1.35] | -0.77 | 0.444 |
|  |  | Adjusted | 0.22 | [-0.51, 0.95] | 0.61 | 0.547 |
| Terminal |  | Unadjusted | 0.17 | [-0.7, 1.05] | 0.40 | 0.692 |
|  |  | Adjusted | 0.10 | [-0.69, 0.9] | 0.26 | 0.795 |
| Full | Monogononta-Ploima | Unadjusted | -0.04 | [-0.98, 0.91] | -0.08 | 0.940 |
|  |  | Adjusted | -0.35 | [-1.3, 0.6] | -0.74 | 0.464 |
| Terminal |  | Unadjusted | -0.38 | [-2.18, 1.42] | -0.42 | 0.677 |
|  |  | Adjusted | -0.44 | [-1.5, 0.63] | -0.82 | 0.416 |
| Full | Female | Unadjusted | -0.43 | [-1.02, 0.16] | -1.47 | 0.147 |
|  |  | Adjusted | -0.43 | [-1.13, 0.27] | -1.22 | 0.227 |
| Terminal |  | Unadjusted | -0.19 | [-0.88, 0.5] | -0.56 | 0.581 |
|  |  | Adjusted | -0.19 | [-0.93, 0.55] | -0.51 | 0.612 |
| Full | Male | Unadjusted | -0.40 | [-0.99, 0.19] | -1.35 | 0.181 |
|  |  | Adjusted | -0.40 | [-1.1, 0.3] | -1.15 | 0.254 |
| Terminal |  | Unadjusted | -0.06 | [-0.74, 0.62] | -0.17 | 0.863 |
|  |  | Adjusted | -0.06 | [-0.79, 0.67] | -0.16 | 0.875 |
| Full | PAC | Unadjusted | 0.23 | [-0.32, 0.78] | 0.85 | 0.399 |
|  |  | Adjusted | 0.22 | [-1.34, 1.78] | 0.28 | 0.779 |
| Terminal |  | Unadjusted | 0.44 | [-0.21, 1.09] | 1.34 | 0.184 |
|  |  | Adjusted | 0.63 | [-0.94, 2.19] | 0.80 | 0.425 |
| Full | Time Lag | | 0.00 | [-0.02, 0.02] | 0.02 | 0.982 |
| Terminal |  |  | 0.00 | [-0.03, 0.02] | -0.32 | 0.751 |
| Full | Publication Bias | | -0.48 | [-1.57, 0.6] | -0.89 | 0.379 |
| Terminal |  |  | -0.30 | [-1.08, 0.48] | -0.77 | 0.442 |

**R code for estimating least-squared regression from published age-specific adult lifespan means and SEs.**

*'dat' is a datafile with rows corresponding to maternal age and columns corresponding to age value, sample size, mean offspring adult lifespan, and the corresponding standard deviation*

colnames(dat) = c("x","n", "y.mean","y.sd")

get.slope <- function(dat){

WSS.optim <- function(par){

SSy = 0

y.hat = par[1] + par[2]*dat$x

for(i in 1:nrow(dat)){

int = .1 #an arbitratily small constant

range = seq(-6 *dat$y.sd[i] + dat$y.mean[i], 6 *dat$y.sd[i] + dat$y.mean[i], by = int)

for (j in range){SSy = SSy + dat$n[i]*dnorm(x = j, mean = dat$y.mean[i], sd=dat$y.sd[i])*int*(j-y.hat[i])^2}}

SSy}

result <- optim(par = c(1,2), fn = WSS.optim)

xbar <- weighted.mean(x = dat$x, w = dat$n)

SSx = 0

for(i in 1:nrow(dat)){SSx = SSx + dat$n[i]*(dat$x[i] - xbar)^2}

num = result$value/(sum(dat$n)-2)

SE = sqrt(num/SSx)

data.frame(estimate = result$par[2], SE = SE)}

get.slope(dat)
